# Customized genomes for human and mouse ribosomal DNA mapping

**DOI:** 10.1101/2022.11.10.514243

**Authors:** Subin S. George, Maxim Pimkin, Vikram R. Paralkar

## Abstract

Ribosomal RNAs (rRNAs) are transcribed from rDNA repeats, the most intensively transcribed loci in the genome. Due to their repetitive nature, there is a lack of genome assemblies suitable for rDNA mapping, creating a vacuum in our understanding of how the most abundant RNA in the cell is regulated. Our recent work ^1^ revealed binding of numerous mammalian transcription and chromatin factors to rDNA. Several of these factors were known to play critical roles in development, tissue function, and malignancy, but their potential rDNA roles had remained unexplored. Our work demonstrated the blind spot into which rDNA has fallen in genetic and epigenetic studies, and highlighted an unmet need for public rDNA-optimized genome assemblies.

We customized five commonly used human and mouse assemblies - hg19 (GRCh37), hg38 (GRCh38), hs1 (T2T-CHM13), mm10 (GRCm38), mm39 (GRCm39) - to render them suitable for rDNA mapping. The standard builds of these genomes contain numerous fragmented or repetitive rDNA loci. We identified and masked all rDNA-like regions, added a single rDNA reference sequence of the appropriate species as a ∼45kb chromosome R, and created annotation files to aid visualization of rDNA features in browser tracks. We validated these customized genomes for mapping of known rDNA binding proteins, and present in this paper a simple workflow for mapping ChIP-seq datasets. These resources make rDNA mapping and visualization readily accessible to a broad audience.

Customized genome assemblies, annotation files, positive and negative control tracks, and Snapgene files of standard rDNA reference sequence are deposited to GitHub.

## INTRODUCTION

Ribosomes are essential ribonucleoproteins composed in eukaryotes of large 60S and small 40S subunits ^2^. RNA components of the ribosome, comprising a majority of its mass, are termed rRNAs (28S, 5.8S, 5S rRNAs in the 60S subunit, and 18S rRNA in the 40S subunit), and collectively form >80% of total cellular RNA. Of these, 18S, 5.8S and 28S rRNAs are produced in mammals through stepwise cleavage of a 13.4 kb 47S pre-rRNA, the transcription of which from rDNA genes by RNA Polymerase I (RNA Pol I) is the rate-limiting step of ribosome biogenesis. rDNA genes, each containing a transcribed region (producing 47S pre-rRNA) followed by a ∼30 kb intergenic spacer, are present in hundreds of near-identical copies (typically 200-600 per cell) in repetitive tandem arrays called Nucleolar Organizing Regions (NORs) ^3^ distributed across 5 chromosomes in humans and mice ^4,5^. rDNA genes are the most intensively transcribed loci in dividing cells, and account for the bulk of all transcription ^6,7^.

The repetitive nature of rDNA has placed substantial bioinformatic barriers in its study. Most standard genome assemblies do not contain intact rDNA sequences, and instead contain a scattering of rDNA fragments, inhibiting the ability of investigators to effectively map high-throughput datasets to them ^1^. The recent telomere-to-telomere (hs1/T2T-CHM13) human genome assembly contains multiple full NOR arrays ^8^, however the presence in this assembly of 219 rDNA repeats creates non-trivial mapping and interpretive complexities. A handful of groups with ribosome biogenesis expertise have mapped datasets to rDNA using privately-generated rDNA-containing assemblies ^9–12^, but, to our knowledge, these assemblies are not publicly deposited. The historical lack of suitable public genomes has led investigators at large to simply ignore rDNA when profiling the binding sites of their transcriptional regulators of interest, creating a significant vacuum in the understanding of how the most abundant RNA in the cell is regulated. Recently published work by our group illustrated the extent of this vacuum, when, through unbiased mapping of ∼2200 ChIP-seq datasets for ∼250 target proteins to rDNA, we identified conserved patterns of rDNA binding for numerous hematopoietic transcription factors and chromatin regulators ^1^. Several of the transcription factors, including the CEBP family, IRF family, and SPI1, all of which have critical roles in hematopoiesis, immune cell development and function, and leukemogenesis, showed consistent, robust peaks at canonical motif sequences. Several of these motifs were conserved between human and mouse. Most of these factors had been extensively studied for decades, and had been subjected to ChIP-seq by dozens of investigators, but their striking rDNA occupancy had passed unnoticed due to a lack of appropriate bioinformatic tools. We further demonstrated that the myeloid transcription factor CEBPA promotes RNA Pol I occupancy on rDNA in myeloid progenitors ^1^. Our work used mammalian hematopoiesis as a model system, but we have no reason to believe that binding of cell-type-specific transcriptional regulators to rDNA is restricted to just one tissue; this is potentially a universal mechanism that is repurposed across all tissues to modulate rRNA transcription rates per tissue-specific needs, but whose prevalence is unknown due to a dearth of suitable mapping tools. Collectively, our work highlighted an unmet need for public assemblies that can be easily used for rDNA mapping.

In this paper, we present human and mouse genome assemblies customized for rDNA mapping (*hg19-rDNA v1*.*0, hg38-rDNA v1*.*0, hs1-rDNA v1*.*0, mm10-rDNA v1*.*0, and mm39-rDNA v1*.*0*). We generated these assemblies by masking degenerate rDNA-matching loci in standard human and mouse genomes, and inserting a new Chromosome R (chrR) composed of a single rDNA reference sequence of the appropriate species. We have made these genomes accessible by depositing them to GitHub, along with annotation files and control mapping tracks. Here, we present validation for the use of these genomes for rDNA ChIP-seq, along with a simple workflow to produce rDNA mapping tracks.

## RESULTS

### Generation of customized human and mouse genomes with rDNA

Human KY962518.1 (44,838 nt) and mouse Genbank BK000964.3 (45,306 nt) reference rDNA sequences are available from NCBI Genbank ^13^, and are used in the literature as consensus reference rDNA sequences for their respective species ^8,14–16^. Each reference contains two promoters (spacer promoter and 47S promoter), a 13.4 kb transcribed region, and intergenic spacer sequence (IGS) (Fig 1A). The human hg19 (GRCh37), hg38 (GRCh38), hs1 (T2T-CHM13), and mouse mm10 (GRCm38), mm39 (GRCm39) genome assemblies each contain multiple partial, fragmented, or repetitive rDNA-matching loci. We identified and masked between 418 to 541 such regions in the genomes (Table S1, see Methods), masking total regions ranging from 252 kb (mm10) to 12 mb (hs1). We then added to these masked genomes a single full-length rDNA sequence of the respective species as an additional chromosome R (chrR) to generate customized genomes (Fig 1B).

**Figure 1.**
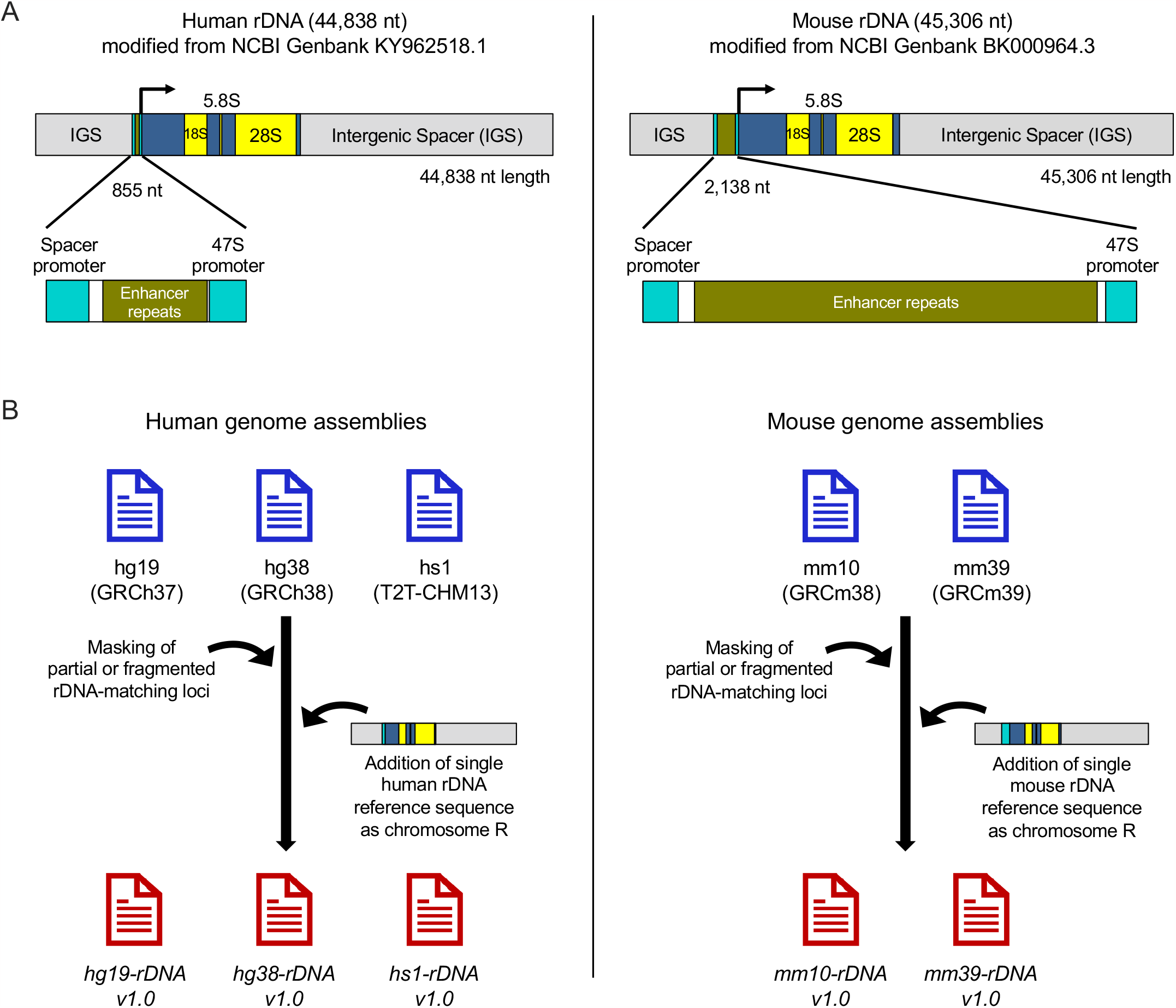
Generation of customized human and mouse genomes with rDNA. (A) Schematic of human and mouse reference rDNA: Reference human (NCBI Genbank KY962518.1) and mouse (NCBI Genbank BK000964.3) rDNA sequences, modified by cutting the original Genbank references at 35,500 and 36,000 nt positions respectively, and transposing the downstream sequence (containing the terminal part of IGS and promoters) upstream of the transcription start site. The transcribed region is depicted in blue and yellow bands, with the sequences corresponding to mature 18S, 5.8S, 28S rRNAs in yellow and those corresponding to external and internal transcribed spacers (ETS and ITS) in blue. The intergenic spacer (IGS) is shown in grey, and promoters are shown in inset detail, with spacer and 47S promoters in cyan. The same color combinations are used to mark rDNA regions in the annotation BED files deposited to GitHub. (B) Generation of custom genomes: Fragmented or repetitive rDNA sequences were masked from human and mouse genome assemblies, and a single reference rDNA sequence of the appropriate species was added on as chromosome R (chrR) to generate the human *hg19-rDNA v1*.*0, hg38-rDNA v1*.*0, hs1-rDNA v1*.*0* genomes, and the mouse *mm10-rDNA v1*.*0, mm39-rDNA v1*.*0* genomes.

### Workflow for mapping to customized genomes with rDNA

Aiming to make routine rDNA mapping accessible to users with no more than basic Linux command line experience, we designed a simple workflow optimized for mapping of high-throughput datasets to the customized genomes (Fig 2). Given the repetitive nature of rDNA, and given that all reads deriving from rDNA will map to the new chrR, even datasets with lower read numbers are capable of giving interpretable chrR coverage, however we recommend using datasets with at least 10 million total reads. We recommend running Trimmomatic ^17^ with standard parameters. We recommend Bowtie2 ^18^ for indexing custom rDNA containing reference genomes and mapping with -X 2000 parameter to permit paired-end sequence mapping with accommodation for 1-2 kb indel variation that has been documented in rDNA repeats ^19^. We recommend filtering SAM files produced by Bowtie2 with Samtools view ^20^ -q 1 to allow for some residual multi-mapping that may occur despite genome masking. After sorting and indexing the resultant BAM files with Samtools ^20^, we recommend using DeepTools BamCoverage ^21^ with -bs 1 to generate nucleotide-resolution tracks. PCR duplicate removal is not recommended; dense overlapping read coverage is expected on chrR due to the presence of hundreds of rDNA copies per cell, leading to the unavoidable presence of independent reads with the same start and end sites, confounding PCR duplicate removal tools. Given that each individual human or mouse has a different number of rDNA copies in its genome, and given that biases arising from library preparation and sequencing protocols can produce some unevenness in background coverage density across the rDNA gene sequence ^22^, users of our customized genomes should anticipate that different cell lines, tissues, and sample preparation methodologies will yield chrR background signal of varying depth and slight idiosyncratic non-uniformity that could be misinterpreted as peak signal if appropriate controls are not used. To robustly distinguish specific signal from background variation, sequencing artifact, or PCR duplication, the inclusion of at least two biological ChIP-Seq replicates, along with paired input and IgG pulldown samples prepared through the same protocol and with the same number of PCR cycles, is critical. We also mention at the bottom of Fig 2 a method of track normalization, to be used if quantitative signal comparison between datasets is required.

**Figure 2.**
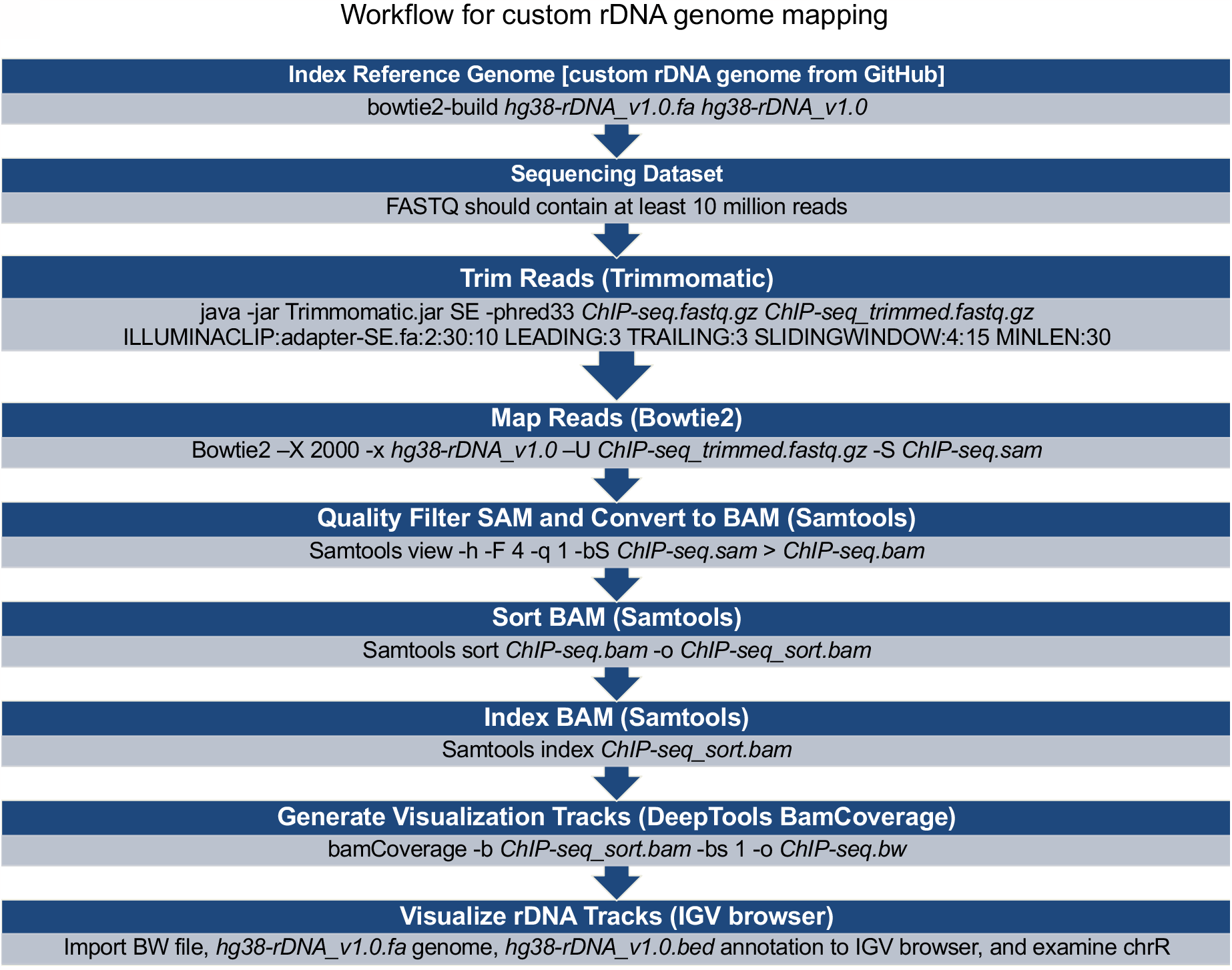
Workflow for mapping to customized genomes with rDNA. Stepwise workflow for mapping of hypothetical high-throughput dataset *ChIP-seq*.*fastq*.*gz* to rDNA, involving sequential steps of adapter trimming with Trimmomatic, mapping with Bowtie2, quality filtering with Samtools view, and generation of visualization tracks with BamCoverage. Additional or optional parameters are provided below the workflow. • <u>Trimmomatic</u>: Use SE or PE for single vs paired-end reads per standard parameters. For increased speed with greater memory usage, may run with ‘-threads 8’ or ‘-threads 16’ parameter. • <u>Bowtie2</u>: For increased speed with greater memory usage, may run with ‘-p 8’ or ‘-p 16’ parameter. • <u>BamCoverage</u>: • Normalization is not required for basic exploratory mapping, but if planning to compare mapping signal between datasets, BamCoverage may be run with the ‘--scaleFactor’ parameter, providing an arbitrary scaling number inversely proportional to the total sequenced reads per dataset. Default scaleFactor is 1. • For rapid track generation restricted to rDNA, may use ‘--region chrR’, which will output only chrR coverage. • May use larger bin size ‘-bs’ for faster speed but lower nucleotide resolution coverage. • <u>PCR duplicates</u>: We do not recommend removing PCR duplicates.

### Validation of customized genomes with rDNA

To validate the ability of our customized genomes and workflow to yield valid rDNA tracks, we picked four proteins whose rDNA ChIP-seq mapping patterns are known ^1,11^: CTCF (Insulator transcription factor), TBP (TATA-binding protein), POLR1A (core catalytic subunit of RNA Pol I), and UBTF (HMG-box protein that binds and activates rDNA). We picked as a negative control EP300, a transcriptional coactivator known to bind promoters and enhancers of Pol II-transcribed genes ^23^ but not known to bind rDNA. Using the public repositories ENCODE ^24^ and GEO/SRAdb ^25^, we identified human and mouse ChIP-seq datasets for each of these factors, and downloaded their raw FASTQ files as well as matched background controls (Input or IgG deposited in the same accession) (links to original datasets are provided in Table S2). We passed these datasets through the workflow in Fig 2, and visualized BigWig (.bw) tracks on the IGV browser ^26^ along with the genome and annotation files (tracks for *hg38-rDNA* and *mm39-rDNA* are shown in Fig 3A, 3B). In all cases, background controls (Input and IgG) showed relatively flat background chrR coverage of significantly lower intensity compared to specific factor pulldown. UBTF and POLR1A showed the expected broad signal throughout the transcribed region, TBP showed the expected peaks at the spacer and 47S promoters, and CTCF showed the expected sharp peak slightly upstream of TBP at the spacer promoter ^1,11^. EP300, as expected, did not show any rDNA occupancy (Fig 3A, 3B). Similar tracks were observed on mapping these datasets to the *hg19-rDNA, hs1-rDNA, mm10-rDNA* genomes (not shown; tracks deposited to GitHub). Thus, we have validated our customized genomes and workflow for high-throughput dataset mapping.

**Figure 3:**
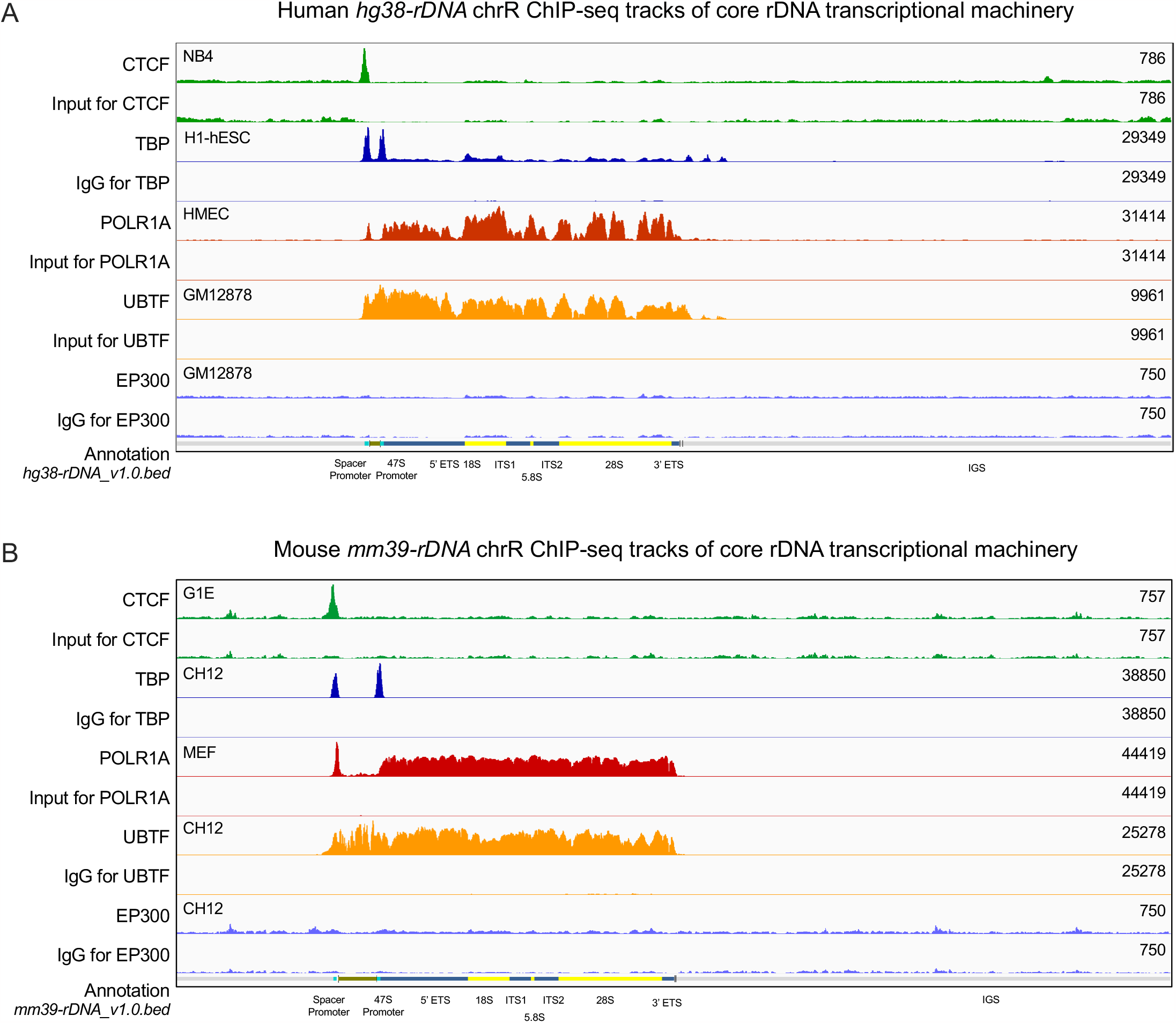
Validation of customized genomes with rDNA. IGV browser track visualizations of: (A) Human (*hg38-rDNA* shown as sample) and (B) Mouse (*mm39-rDNA* shown as sample) chrR ChIP-seq mapping. Known rDNA binding proteins CTCF, TBP, POLR1A, UBTF are shown as positive controls, EP300 is shown as a negative control, and matched Input or IgG are shown as background controls. Factor names are listed on the left, cell types for each ChIP-Seq are shown on the top left of track pairs, and Y-axis scale of each track is listed on the right. Matched pairs of control and background tracks are set to the same Y-axis scale. At the base of each image are rDNA segment annotations. 5’ ETS and 3’ ETS are External Transcribed Spacers, ITS1 and ITS2 are Internal Transcribed Spacers. IGS is the Intergenic Spacer.

### Accessibility of customized genomes and annotation files

All customized human and mouse genomes and associated files have been deposited to GitHub (https://github.com/vikramparalkar/rDNA-Mapping-Genomes). Specifically, FASTA files for 3 human (*hg19-rDNA v1*.*0, hg38-rDNA v1*.*0, hs1-rDNA v1*.*0)* and 2 mouse *(mm10-rDNA v1*.*0, mm39-rDNA v1*.*0*) genomes have been deposited, along with annotation BED files (containing standard RefSeq gene annotations plus new chrR annotations), and Bigwig files for positive and negative controls tracks for each genome.

## DISCUSSION

The public availability of genome assemblies ^27^, enabling widespread profiling in diverse cell types of chromatin accessibility, histone modifications, transcription factor binding, and chromosome conformation, has facilitated unprecedented progress over the past two decades in our understanding of gene regulation in health and disease. However, certain regions of the genome remain poorly studied due to their repetitive nature ^28^. Ribosomal DNA genes, organized in tandem arrays in NORs, are a prominent example. Due to the inability of standard genome assemblies and mapping pipelines to yield interpretable rDNA data, a stark contrast has emerged between the depth of our understanding of coding genes compared to rDNA genes. Given that rRNA is the most abundant cellular RNA, and accounts for the bulk of all transcription, this is a significant bioinformatic blind spot.

We attempt in this paper to address this deficiency by generating and depositing to GitHub customized versions of multiple commonly used human and mouse genomes containing a single copy of reference rDNA as an added chromosome, along with annotation files, sample tracks, and a simple mapping workflow. This method can be easily extended to any new assembly that may be released, as well as to other species for which full-length reference rDNA sequences are available. Our recent work ^1^ indicates that rDNA is bound and regulated in complex eukaryotes by a variety of cell-type-specific transcription factors. The implications of such binding for evolution, cell biology, development, and oncogenesis are substantial, and warrant a systematic exploration of whether modulation of ribosome production rates through control of rRNA transcription is a conserved function of many transcription factors and chromatin regulators, and whether this regulation is hijacked in malignancies to promote rapid ribosome biogenesis. Identification of such factors will only become possible with the routine incorporation of rDNA-containing genomes into mapping pipelines.

rDNA repeats, though fairly uniform in their tandem repeat arrangements, are not perfectly identical, and show variations in the form of single nucleotide polymorphisms scattered throughout the repeat, as well as small insertions and deletions in the IGS ^16,19,29,30^. These variations can be seen between repeats within the same cell as well as between individuals within the same species. The next challenge for rDNA bioinformatics will be to accommodate rDNA variants. Continued work by the T2T Consortium ^8^ and the Human Pangenome Project ^31^ will, over time, reveal the full extent of variation in rDNA number and sequence. We anticipate that future standard genome assemblies of all species will contain a systematized diversity of rDNA repeat sequences, and the next generation of mapping tools will need to grapple with the peculiar challenges of effectively mapping to these repetitive loci.

In summary, we provide in this paper customized genomes, annotation files, and a simple workflow to make human and mouse rDNA mapping accessible to non-specialists in ribosome biogenesis. We envision that routine integration of these genomes into standard bioinformatic pipelines will allow the discovery and characterization of novel regulators of rDNA relevant to normal and disease biology.

## EXPERIMENTAL PROCEDURES

### Identifying regions of human and mouse genome assemblies with sequence similarity to reference rDNA

The full length rDNA sequence is ∼45 kilobases (kb) in human and mouse ^19,32^, and all human and mouse assemblies contain numerous partial, fragmented, or repeated rDNA sequence stretches. To systematically identify these regions for masking, we used four methods: (i) We obtained from the USCS genome browser ^33^ a list of all Repeatmasker ^34^ annotations titled ‘LSU-rRNA’ or ‘SSU-rRNA’ (large and small subunit rRNA). (ii) We obtained from NCBI Genbank ^13^ the reference sequences for human and mouse rDNA (KY962518.1 ^19^ and BK000964.3 ^32^ respectively), and used BLAT ^35^ to identify genomic intervals with matches to the reference sequences. (iii) Modifying a previously reported approach ^22^, we fragmented the KY962518.1 and BK000964.3 rDNA gene reference sequences to a collection of 50 nt “reads” that we mapped to genomes of the respective species using Bowtie2 ^18^ with the -k 10 parameter to include multiple mapped locations. We then parsed the output SAM file to identify stretches of the genome with consecutive mapping of at least 8 of the fragmented “reads”, including those in which adjacent stretches were separated by less than 200 nt. (iv) For the hs1 (T2T-CHM13) genome, which contains full NOR arrays on 5 chromosomes, we obtained from the USCS genome browser ^33^ a list of 5 Centromeric Satellite Annotations titled ‘rDNA’. To add buffer regions, we extended intervals from (i), (ii), and (iv) by 100 nt upstream and downstream, and those from (iii) by 500 nt. Because several intervals were overlapping or redundant, we used the Merge tool in the public Galaxy server usegalaxy.org ^36^ to merge the interval groups and produce a final non-redundant set of intervals (Table S1): hg19 (GRCh37): We finalized for masking 473 intervals ranging from 225 nt to 61 kb in length, with mean and median lengths of 763 nt and 331 nt respectively. One interval was longer than 45 kb (the reference rDNA length), and on manual curation were found to be composed of more than one rDNA copy.

hg38 (GRCh38): We finalized for masking 473 intervals ranging from 226 nt to 271 kb in length, with mean and median lengths of 1,862 nt and 323 nt respectively. Four intervals were longer than 45 kb (the reference rDNA length), and on manual curation were found to be composed of sequence combinations corresponding to more than one rDNA copy.

hs1 (T2T-CHM13): We finalized for masking 418 intervals ranging from 1,132 nt to 3.5 mb in length, with mean and median lengths of 29 kb and 5 kb respectively. These included 5 full NOR arrays on chromosomes 13, 14, 15, 21, 22, ranging from 718 kb to 3.5 mb and encompassing 219 rDNA repeats.

mm10 (GRCm38): We finalized for masking 541 intervals ranging from 231 nt to 17 kb in length, with mean and median lengths of 467 nt and 311 nt respectively.

mm39 (GRCm39): We finalized for masking 516 intervals ranging from 223 nt to 32 kb in length, with mean and median lengths of 554 nt and 318 nt respectively.

### Generating masked genome assemblies

We used the bedtools maskfasta ^37^ tool to mask human and mouse rDNA-like loci, and masked a total of 361 kb, 881 kb, 12 mb, 252 kb, and 286 kb in the hg19, hg38, hs1, mm10 and mm39 genomes respectively. We remapped the 50 nt fragmented reads generated from human and mouse rDNA references (see section above) to the masked genomes of the respective species, and found no consecutive mapping of more than 8 reads, confirming adequate masking.

### Adding reference rDNA sequence to masked genome assemblies

The human KY962518.1 (44,838 nt) and mouse BK000964.3 (45,306 nt) reference rDNA sequences both start with the rRNA transcription start site (TSS) and end with the Pol I promoter. Given that these ∼45 kb units are repeated in tandem arrays in NORs, their 3’ ends can be seen as continuous with their 5’ ends, and we therefore modified their sequences to bring their promoters upstream of their TSSs for ease of track visualization. We did so by cutting the human KY962518.1 sequence at the 35,500 nt position and the mouse BK000964.3 sequence at the 36,000 nt position, and transposed the sequences downstream of the cut to upstream of the TSSs, such that in the modified orientation, the rDNA sequences began with ∼10 kb of IGS before transitioning into promoters and transcribed regions (final orientation shown in Fig 1A). We chose the cut sites based on our rDNA mapping study of ∼2200 ChIP-seq datasets ^1^, which did not show overlap of these sites with any transcription factor, chromatin protein, or histone modification peaks. We then added the modified rDNA reference sequences to masked genomes of the respective species as an additional chromosome R (chrR) (Fig 1B). These customized assemblies are *hg19-rDNA v1*.*0, hg38-rDNA v1*.*0, hs1-rDNA v1*.*0, mm10-rDNA v1*.*0, mm39-rDNA v1*.*0*.

### Generating annotation files containing chromosome R

We obtained from the USCS genome browser ^33^ the RefGene ^38^ annotation BED files for each assembly, swapped RefSeq NM/NR IDs with Gene Name IDs, and added chrR annotations depicting the known locations of spacer and 47S promoters, rRNA transcript segments, and TTF1 binding sites within the rDNA references ^1,11,39^. The final annotation BED files were confirmed to be compatible with the IGV browser ^26^ for visualization of chrR as well as for genome wide visualization of standard gene structures.

## DATA AVAILABILITY

Customized genome assemblies, annotation files, and positive and negative control mapping tracks are available through GitHub at: https://github.com/vikramparalkar/rDNA-Mapping-Genomes

## ACKNOWLEDGEMENTS

V.R.P is supported by the National Institute of General Medical Sciences (NIGMS) of the National Institutes of Health (NIH) under award number R35-GM138035.

## AUTHOR CONTRIBUTIONS

**Vikram R Paralkar:** Conceptualization, Methodology, Supervision, Funding acquisition, Project administration, Writing - Original Draft. **Subin S George:** Methodology, Software, Validation, Investigation, Data Curation, Visualization, Writing - Original Draft. **Maxim Pimkin:** Conceptualization, Writing - Review & Editing

